# The CD4 T cell epigenetic JUNB+ state is associated with proliferation and exhaustion

**DOI:** 10.1101/2024.01.05.573875

**Authors:** Ionut Sebastian Mihai, Martin Selinger, Nicole Boucheron, Mattias Forsell, Isabelle Magalhaes, Johan Trygg, Johan Henriksson

**Author notes:** Shared first author (Ionut Sebastian Mihai, Martin Selinger). **Contact:** Ionut Sebastian Mihai, https://orcid.org/0000-0002-9322-5879 Martin Selinger, https://orcid.org/0000-0002-5420-9702 Nicole Boucheron, https://orcid.org/0000-0002-4979-8311 Mattias Forsell, https://orcid.org/0000-0001-6904-742X Isabelle Magalhaes, https://orcid.org/0000-0003-0440-6924 Johan Trygg, http://orcid.org/0000-0003-3799-6094 Johan Henriksson, https://orcid.org/0000-0002-7745-2844 or.

## Abstract

Adoptive cell therapy (ACT) requires the in vitro expansion of T cells, a process where currently several variables are poorly controlled. As the state and quality of the cells affects the treatment outcome, the lack of insight is problematic. To get a better understanding of the production process and its degrees of freedom, we have generated a multiome CD4 T cell single-cell atlas. We find in particular a JUNB+ epigenetic state, orthogonal to traditional CD4 T cell subtype categorization. This new state is present but overlooked in previous transcriptomic CD4 T cell atlases. We characterize it to be highly proliferative, having condensed and actively remodeled chromatin, and correlating with exhaustion. JUNB+ subsets are also linked to memory formation, as well as circadian rhythm, connecting several important processes into one state. To dissect JUNB regulation, we also derived a gene regulatory network (GRN) and developed a new explainable machine learning package, Nando. We propose potential upstream drivers of JUNB, verified by other atlases and orthogonal data. We expect our results to be relevant for optimizing in vitro ACT conditions as well as modulation of gene expression through novel gene editing.

## Introduction

CD4+ T cells are key components of the adaptive immune system which carry out a wide variety of actions: from recruiting and activating other cellular components of the immune system to the site of infection, to fine-tuning the immune response. Because of their ability to target particular cells (such as cancer cells) via the T cell receptor, T cells are used for adoptive cell therapy (ACT). However, many T cell subtypes exist, and their composition is crucial depending on their application. The compositional choice is nontrivial, and the use of “pure” subtypes is not always beneficial; e.g. mixed CD4 and CD8 T chimeric antigen receptor (CAR) T cells are better against glioblastoma^1^. CD4 T cells in turn consist of many subtypes, and ACT with CD4 T regulatory helper cells (Tregs) have been used to avoid graft-versus-host disease^2^, while 4th generation CAR T cells biased towards T helper type 1 (Th1) have been used for cancer therapy^3^. To aid the compositional decision, it is necessary to first understand what T cell states (or types) might be present, starting from the bioreactor where they are produced^4,2^. We have thus generated an atlas of representative CD4 T cells using modern single-cell multiomics technology.

We find both classical and non-classical T cell subtypes, including subtypes not classically expected given the used cytokine cocktails. We also further elaborate on the entanglement between differentiation and proliferation^5^. The Tfh state is intertwined with JUNB, and we find by comparison with other atlases that a new cell type (or state) can be defined as JUNB+. It is generated during most standard differentiation conditions but possibly more linked to Th17. It is associated with proliferation, and expresses several chromatin remodeling factors which are top hits in CRISPR screening for T cell exhaustion. These factors appear crucial for the nature of the JUNB cells, as they have a closed chromatin state and elevated expression of the insulator CTCF. While it is not yet clear how to think about these so-far overlooked JUNB cells (representation of states being non-trivial in general^6^), we show that it has a number of properties relevant for ACT, thus being of high interest.

To further understand how the T cell subtypes are regulated, we have developed a new explainable AI (XAI) approach to dissect how the gene regulatory network (GRN) is wired for each cell state. This makes particular use of our dataset being RNA+ATAC-seq^7^. By comparing our data to other atlases, we find for example ZNF331 to be a potential regulator of the JUNB state. Knowledge of these regulators and their function will inform development of future ACT, e.g., when using genetically reprogrammed CAR T cells. Our atlas and the putative network can be browsed at http://data.henlab.org/.

### Multimodal analysis provides two complementary perspectives of T cell states

To obtain an overview of possible CD4 T helper states during *in vitro* production, isolated naive CD4 T cells from 4 human donors were activated using anti-CD3/CD28 and simultaneously exposed to cytokine cocktails commonly used for differentiation into Th1-, Th2-, Th17-, and Treg subtypes (see methods, overall strategy in Figure 1a). The cells were grown in defined media, without FBS, to adhere to GMP (good manufacturing practice). After 5 days, cells were analyzed for the expression of Th1/Th2/Th17/Treg canonical markers TBX21, GATA3, RORC, and FOXP3, respectively. In all cases, the population of differentiated cells showed increased levels of the respective canonical marker when compared to naïve non-activated cells (Figure S1a-b). Cells from different stimuli were pooled, enriched for viable cells using FACS, and libraries were constructed using 10x Genomics’ single-cell multiomics (RNA+ATAC).

**Figure 1:**
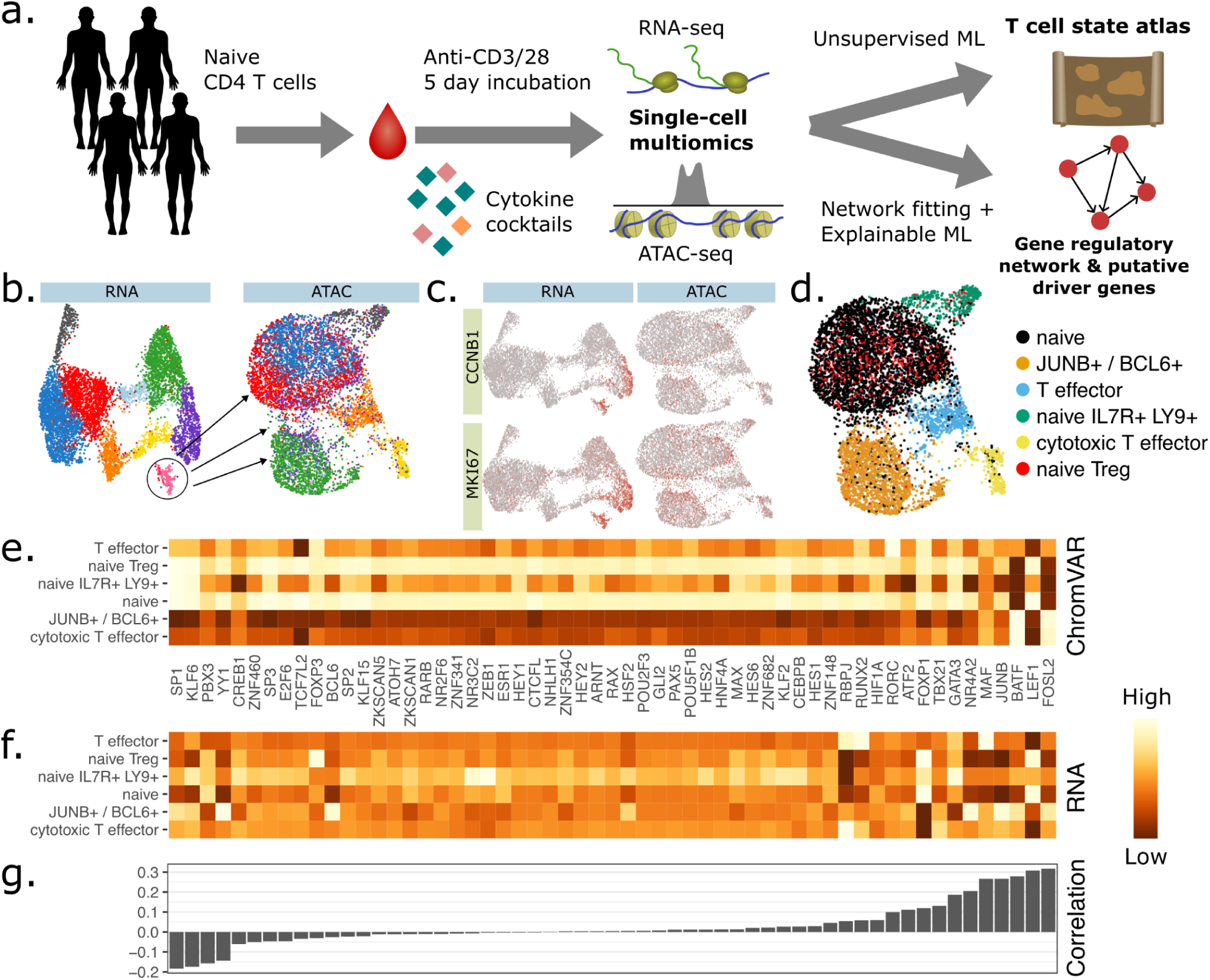
A multiome atlas of in vitro activated CD4 T cells. (**a**) The experimental setup and study design, resulting in a multiome atlas, and a network of predicted driver genes. (**b**) Clustering based on RNA-seq similarity, on RNA-seq and ATAC-seq dimensional reductions. An additional cell cycle cluster is removed in the ATAC-seq view. (**c**) UMAPs for cell cycle linked genes MKI67 and CCNB1, showing how they mainly drive the RNA UMAP. (**d**) The clustering used for the remainder of this study, mainly based on ATAC-seq to avoid cell cycle influence. (**e**) Gene expression of some representative marker genes for each cell type, with (**f**) corresponding motif activities and (**g**) their correlation.

**Supplemental figure 1:**
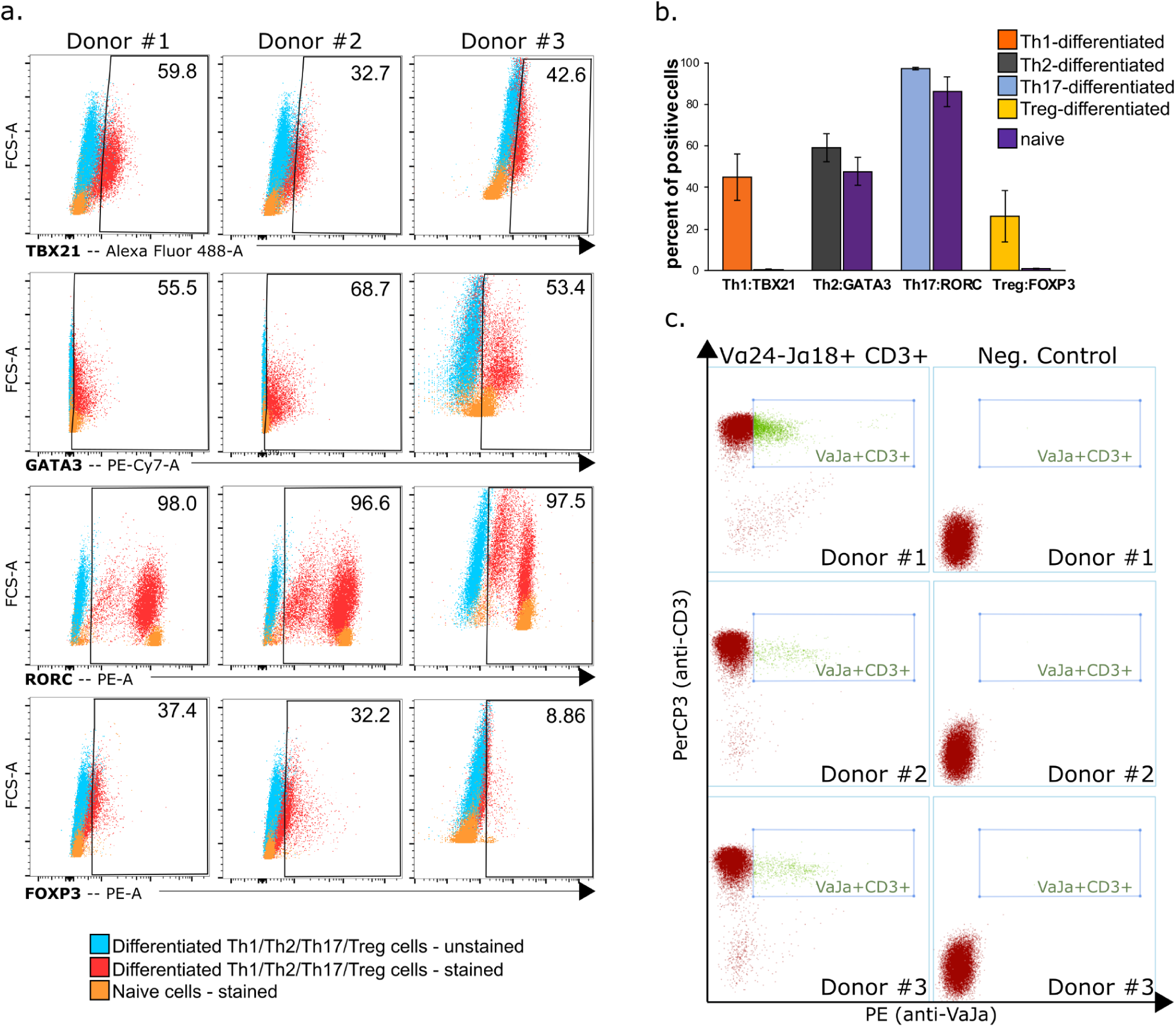
Validation of conditions. (**a-b**) FACS of canonical marker genes show their enrichment in respective conditions. **(c)** FACS show the presence of NKT cells directly after bead-based negative selection.

**Supplemental figure 2:**
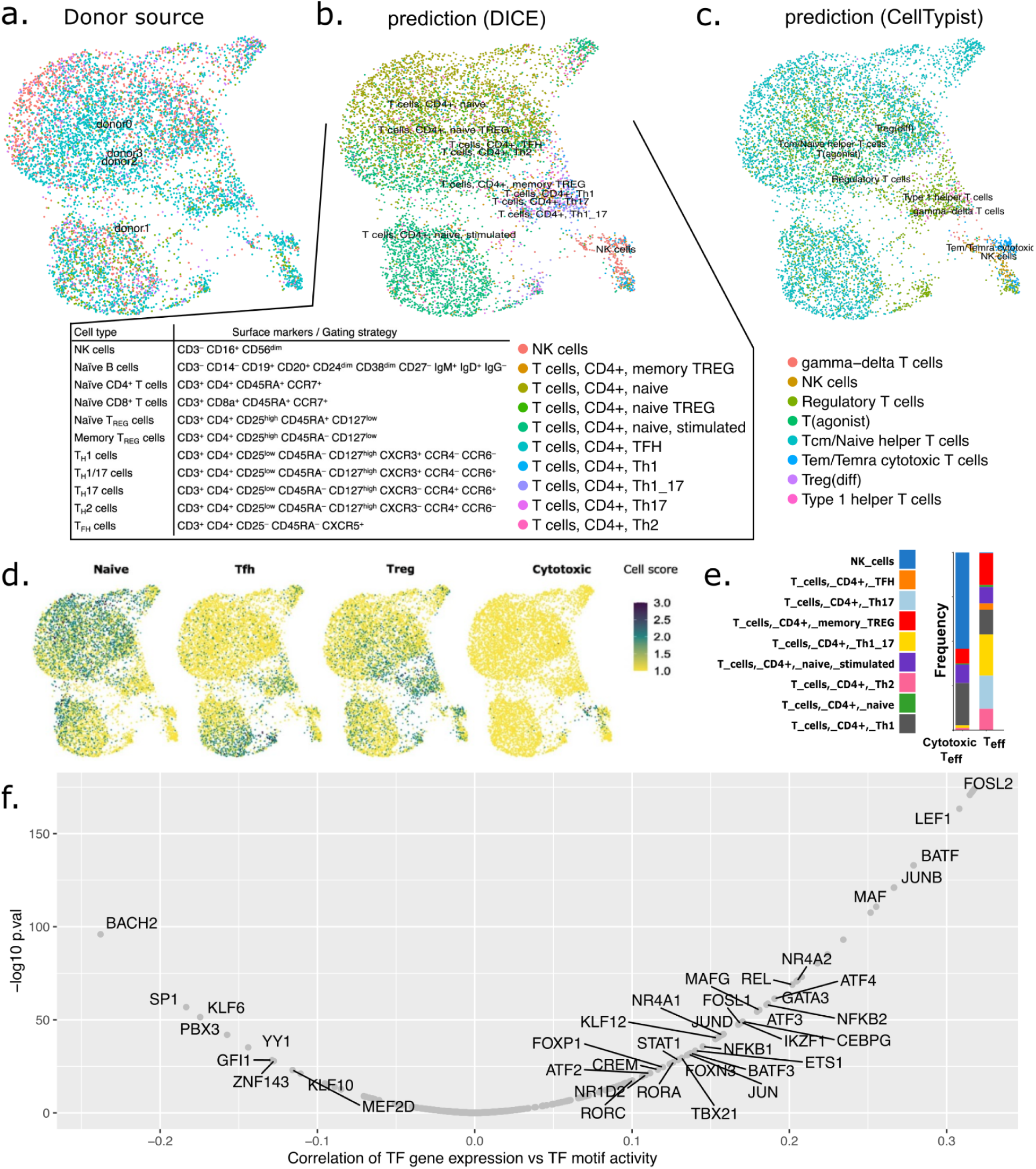
Annotation of cell types. Labels on UMAPs indicate the center position of all cells. **(a**) Cells from different donors mix well, showing that clusters are not specific to a donor. **(b)** Automatic cell type annotation based Schmiedel2022^8^ dataset. **(c)** Automatic cell type annotation using CellTypist. **(d)** Scoring of cell type identity using lists of cell type specific genes. **(e)** Stacked barplot of DICE-annotated subtypes within cytotoxic Teffector and T effector clusters. **(f)** Correlation of ChromVAR motif activity to the expression of the corresponding gene. Where there is high correlation, the regulation is likely mainly due to the transcriptional level of the TF as opposed to posttranslational modifications. All correlations are provided in Supplemental file S9.

Using UMAP dimensional reduction and Leiden clustering, we found that RNA-seq and ATAC-seq provide two similar but complementary perspectives of the CD4 T cell composition (Figure 1b). In particular, cell cycle genes affect the RNA-seq based clustering/UMAP more than for ATAC-seq. This can be seen by common cell cycle marker genes such as MKI67 or CCNB1 (Figure 1c). Cells closer to mitosis (e.g. MKI67^hi^, CCNB1^hi^) tend to land in between the large clusters in ATAC-seq, or along their borders. The RNA-seq-based clustering instead tends to co-cluster all MKI67^hi^ and CCNB1^hi^ cells together. This suggests that cell cycle is mainly observed at the transcriptional layer. We thus mainly used ATAC-seq to delineate the cell types (Figure 1d, procedure described in Methods). The cell types largely correspond to expectations. Searching for potential impurities that could be of issue for ACT, we only find about 5% NKT cells (Figure S1c). Although CAR NK is gaining interest^9^, we deemed this population too small for deeper characterization. Given the need for speed and scalability for ACT, bead-based negative selection may overall be a good initial choice.

While previous T cell atlases have primarily focused on gene expression, our multimodal readout further enables the analysis of upstream regulating TFs. The difference in the UMAPs of RNA and ATAC suggests that these perspectives are not completely aligned (e.g., not equally driven by cell cycle, but likely also other aspects). To compare these views further, we used ChromVAR to detect TF motifs that are overrepresented in open chromatin regions^10^ (motif activity). Since we performed RNA and ATAC-seq on the same cell, we were able to find TFs correlating expression-wise with motif activity. By the simplifying assumption that the highest correlating TF correlating with a motif’s activity is also the TF binding, we are able to overcome that some TFs share motifs (Figure S2e). This does not rule out that other TFs may also interact, but helps focus on the likely most important TFs. We show the representative marker genes for each cell type (Figure 1e), its corresponding motif (Figure 1f) and their correlation (Figure 1g). Most TFs are positively correlated, meaning that increased expression results in its motifs being more accessible. FOSL2 and LEF1 are the most correlating genes, while the repressor BACH2 is the most negatively correlating (Figure S2e). Both LEF1^11^ and BACH2^12^ regulate stemness, and the high correlation likely reflects their broad activity (since TFs active in many enhancers get higher ChromVAR score). Other TFs are correlated, but much less, showing that the epigenetic and transcriptional views are similar, but not the same. Our atlas instead suggests that these TFs may be post-translationally controlled, and can thus be difficult to reprogram for ACT purposes if other players in fact control their activity. The metabolic regulator HIF1A, upregulated in the hypoxic TME^13^ (tumor microenviroment), is however positively correlated. While it is well-known to be regulated post-translationally by, e.g., PHD1, this is clearly not the only mechanism. Our epigenetically informed atlas thus helps pinpoint TFs that likely can be modulated through overexpression in 4th generation CAR T cells, which carry transgenic payloads^14^.

### JUNB and BCL6 delineate a distinct and hyperproliferative Tfh-like program

We were intrigued by a cluster which we struggled to fit with classical annotation of cell types. It appears primed toward Tfh based on its high expression of BCL6 (Figure 2a). However, it lacks other canonical Tfh markers, such as CXCR5. The transcription factor JUNB is also a good marker of this subset, and has among the highest correlation of motif activity *vs* RNA (Figure S1f). Thus we assumed JUNB to be the more relevant driver gene and decided to focus on it for the remainder of our study.

**Figure 2:**
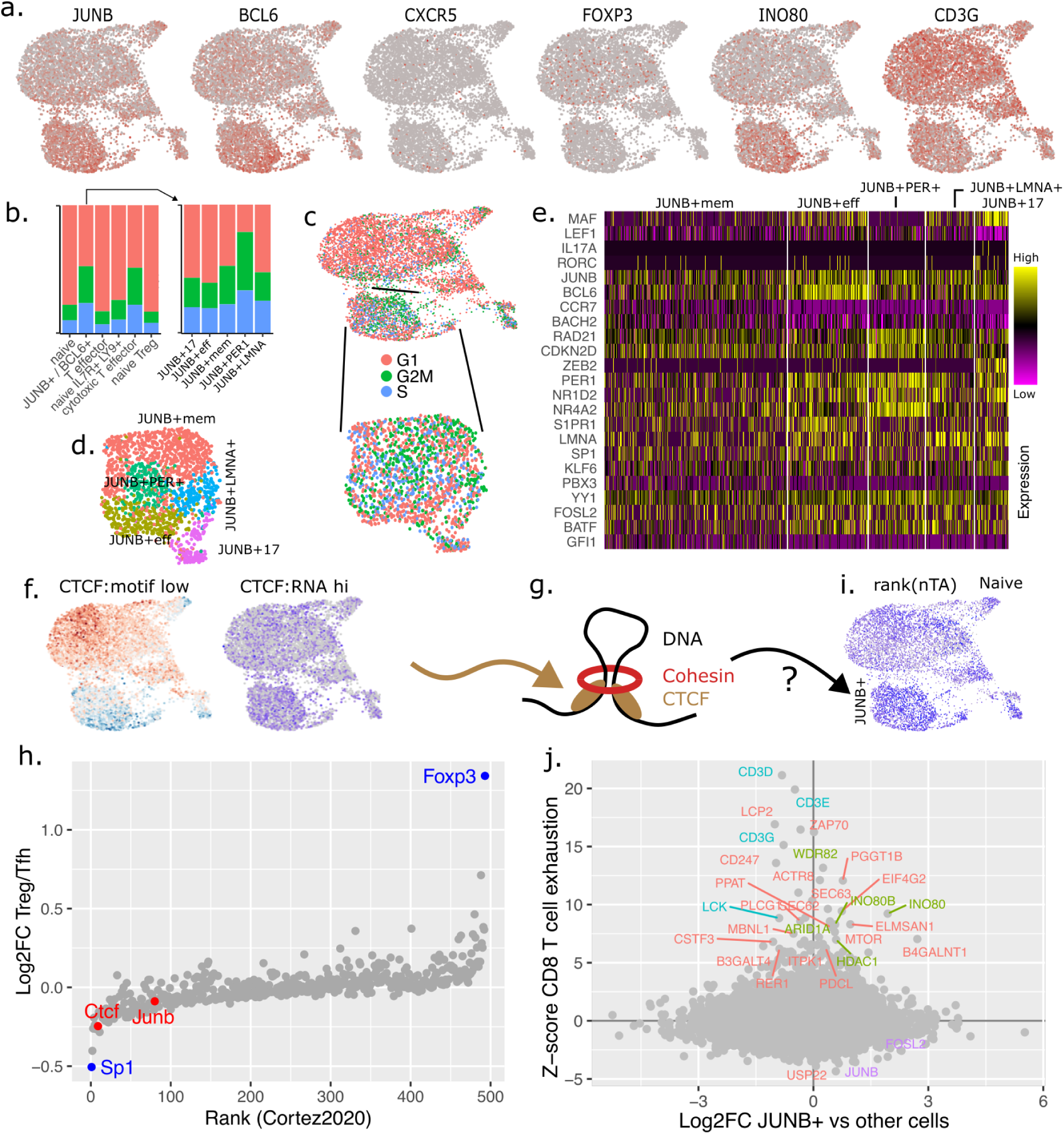
Detailed analysis of the JUNB/BCL6+ cluster. (**a**) Detailed expression patterns on UMAP, showing the gene specificity. (**b)** Cell cycle phase per clusters, shown as fractions and **(c)** on UMAP, for all cells and JUNB+ reclustered cells. **(d)** UMAP of JUNB+ cells, dimensionally reduced separately. **(e)** Heatmap of key genes across JUNB+ subclusters. **(f)** CTCF motif activity and gene expression. **(g)** Representation of a transcriptionally activate domain (TAD) model with CTCF, which guides the cohesin loops and thus 3D structure. **(h)** Detected regulators of Foxp3, from a murine Treg CRISPR screen dataset (Cortez2020)^15^. CTCF and JUNB act antagonistically to FOXP3, which is low in the JUNB state. **(i)** Telomemore analysis predicts that Tfh cells have increased chromatin condensation. **(j)** Detected regulators of exhaustion in CD8 T cells in a CRISPR screen^16^ vs DE genes in our JUNB cluster. Top scoring chromatin remodelling factors are shown in green, and TCR-related proteins in purple. JUNB is among the highest scoring genes for exhaustion.

**Supplemental Figure 3:**
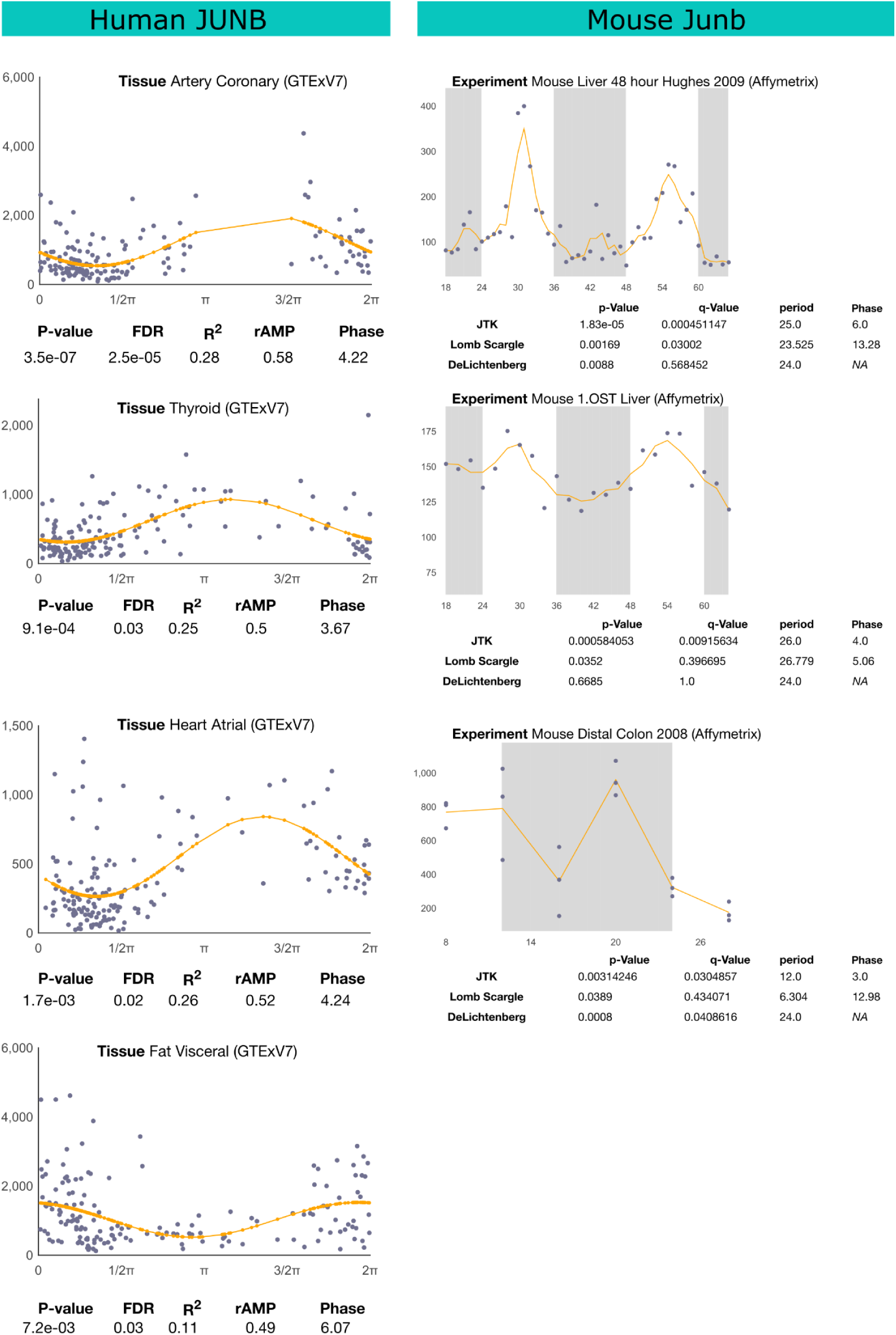
Analysis of JUNB circadian expression across tissues. Reproduced from http://circadb.hogeneschlab.org/, where default search queries were used to find datasets where JUNB is significant^17^.

Based on the gene expression data, we note that not all cells proliferate equally (Figures 2b-c). A new clustering was performed on JUNB+ cells alone to minimize the cell cycle effect (Figures 2d-e). Since the canonical marker BCL6 itself is not cell cycle linked^18^, our interpretation is rather that JUNB+/Tfh is a distinct state that proliferates at a higher rate than other cell types. A small subset of cells (Figure 2e, JUNB+17) express Th17 (IL17A, RORC) and effector (MAF+LEF1-) markers. Th17 has also previously been seen to be hyperproliferative^19^. However, while many marker genes follow the expected cell cycle annotation in the JUNB+ cluster, other genes are expressed in a more cell type dependent manner (Figure 2e). This includes, e.g, RAD21 (part of cohesin) and CDKN2D, and may thus be present to continuously maintain proliferation.

Tfh has been shown to be highly plastic, with cytokine producing subtypes Tfh_1,2,17_ and Tfr have been suggested^20^. Tfh_17_ in particular has a high capacity for self-renewal and plasticity^19^. The UMAP topology and high level of proliferation among our JUNB+/Tfh cells (and previous Tfh_em_ bulk RNA-seq^21^) points to the possibility that Tfh and Th17 might be more similar cell states than previously appreciated, at least under our *in vitro* conditions. Tfh_17_ has been shown to be highly dependent on HIF1A, which is the most expressed at Tfh_em_ and at the entry to Tfh_17_^19^.The Tfh_17_ subcluster is high in ZEB2, a gene highly expressed in T_em_ cells.

Another subcluster is JUNB+PER1+, which possibly has the highest turnover (Figures 2b-c). We are unable to link it to a classical CD4 T cell subtype, but some genes are related to circadian rhythm, including PER1, NR1D2 (Rev-Erbβ) and NR4A2, and might relate to the circadian migration of T cells from lymph nodes^22^. In particular, the migration depends on the oscillation of S1PR1^22^, whose expression is restricted to Tfh, but especially high in Tfh_eff. Thus exit might occur in an effector-like state, possibly analogous with how the murine circadian Rora also appears linked to T cell activation^23^ (however, in our atlas, RORA is mainly expressed in Teff). Interestingly, differentiation of Th_17_ (clustered nearby) is also controlled in a circadian manner^24^. JUNB itself is circadian in some tissues (Figure S3), but we have not found evidence for this in CD4 T cells. Since success of surgery is affected by when during the day it is performed^25^, further insight into this cluster might also be beneficial for ACT.

To summarize, our atlas contains cells similar to the naive Tconv, naive Treg, T effector, cytotoxic T effector and Tfh subtypes. Noteworthy, we did not aim to polarize any cells toward Tfh, but a similar JUNB+ state seems naturally present in the heterogeneous culture. Because efficient *in vitro* cell expansion is key to ACT, and this subset appears somehow linked to proliferation, we decided to characterize it deeper.

### JUNB delineates an epigenetically condensed state, linked to persistence and chromatin remodelling

For the JUNB+ cluster, ChromVAR showed differential CTCF expression/activity (Figure 2f). CTCF is considered a special insulating factor, being present at transcriptionally active domain (TAD) boundaries (Figure 2g). A murine screen has been performed to identify regulators of Treg *vs* Tfh fate (Cortez2020)^15^. Interestingly, CTCF is one of the top regulators of FOXP3 expression (Figure 2h), which is down in JUNB+ (Figure 2a). To get an idea of the overall chromatin state, we analyzed the ATAC-seq data using Telomemore, based on the principle that high ATAC telomere abundance frequently indicates low general accessibility^26^. It appears that the chromatin is particularly and continuously condensed in both JUNB/Tfh and JUNB_17_, (Figure 2i), and it is possibly due to a higher order change in TAD structure, with increased insulation, driven by CTCF and other factors. Telomemore also predicts high condensation of naive T_conv_ and naive Tregs, in line with previous time course ATAC-seq data^5,26^. The large difference in chromatin accessibility in JUNB+ *vs* other clusters opens questions about the nature of this state. T cell plasticity has previously been shown using lineage tracing^27^. The gap between JUNB+ and other clusters suggest a possibly rapid change of epigenetic state during repolarization. Furthermore, the most similar cells along the cluster boundaries are mitotic (CCNB1, Figure 1d), supporting the idea that polarization may be coupled to a cell division and simultaneous large chromatin rearrangement. The chromatin remodeler INO80 is high in JUNB+, and having been shown to repress and license bivalent genes^28^, offering one possible mechanism for plasticity.

To see how our cell states relate to T cell exhaustion, a key problem in ACT, we compared our data to available CRISPR screens. A screen has been performed in murine CD8 (but not CD4) T cells^16^, showing JUNB as a top regulator. The coexpressed FOSL2 is also a top regulator. Interestingly, overlapping the top hits shows that a large fraction of them are differentially expressed in the JUNB cluster *vs* all other clusters, suggesting that the JUNB+ cluster may be crucial for the exhaustion process (Figure 2j). We can only speculate why, but the screen highlights that several top genes are chromatin remodeling factors, such as INO80, HDAC1, ARID1A and WDR82. Because INO80 licenses bivalent genes^28^, and fate in stem cells^28^, exhaustion might rather be an epigenetic process, linked to the plasticity toward other cell states. An alternative or complementary explanation for the exhaustion could also be related to the TCR. Interestingly, the murine exhaustion screen also pinpointed the TCR subunits CD3G (Figure 2j) and CD3D, as well as the related LCK. While this should be expected, we also note that these are all somewhat downregulated in JUNB+ (Figure 2a), and might result in a reduced activation potential of the JUNB+ state. We have previously shown that exhaustion could be decreased by modulating the CAR activation^29^. It is thus possible that the downregulation of TCR genes could be an innate compensatory mechanism to avoid exhaustion, that could be harnessed for ACT. Overall, the JUNB+ state appears riddled with factors related to exhaustion, and a better understanding of this state is likely to result in improved therapy.

### The JUNB+ state is universal and memory associated

To validate that the JUNB state is universally present, we compared our data to other atlases. We found a candidate JUNB+ cluster in naive CD4 T cells, 4h post-activation (Schmiedel2022^8^). These cells have been gated by FACS, enabling us to conclude that it is composed of cells from all canonical subsets as defined by FACS (Figure 3a, S1b). However, this cluster is not enriched in BCL6 (Figure 3b), showing that BCL6 and JUNB are not always co-expressed. Oddly, Schmiedel2022 also contains Tfh cells defined by FACS on CXCR5, but these are not particularly high in either JUNB or BCL6. This suggests that our view of BCL6 and human Tfh may need to be revised. As in our data, the JUNB+ state cannot be explained simply as a cell cycle stage (Figure 3c).

**Figure 3:**
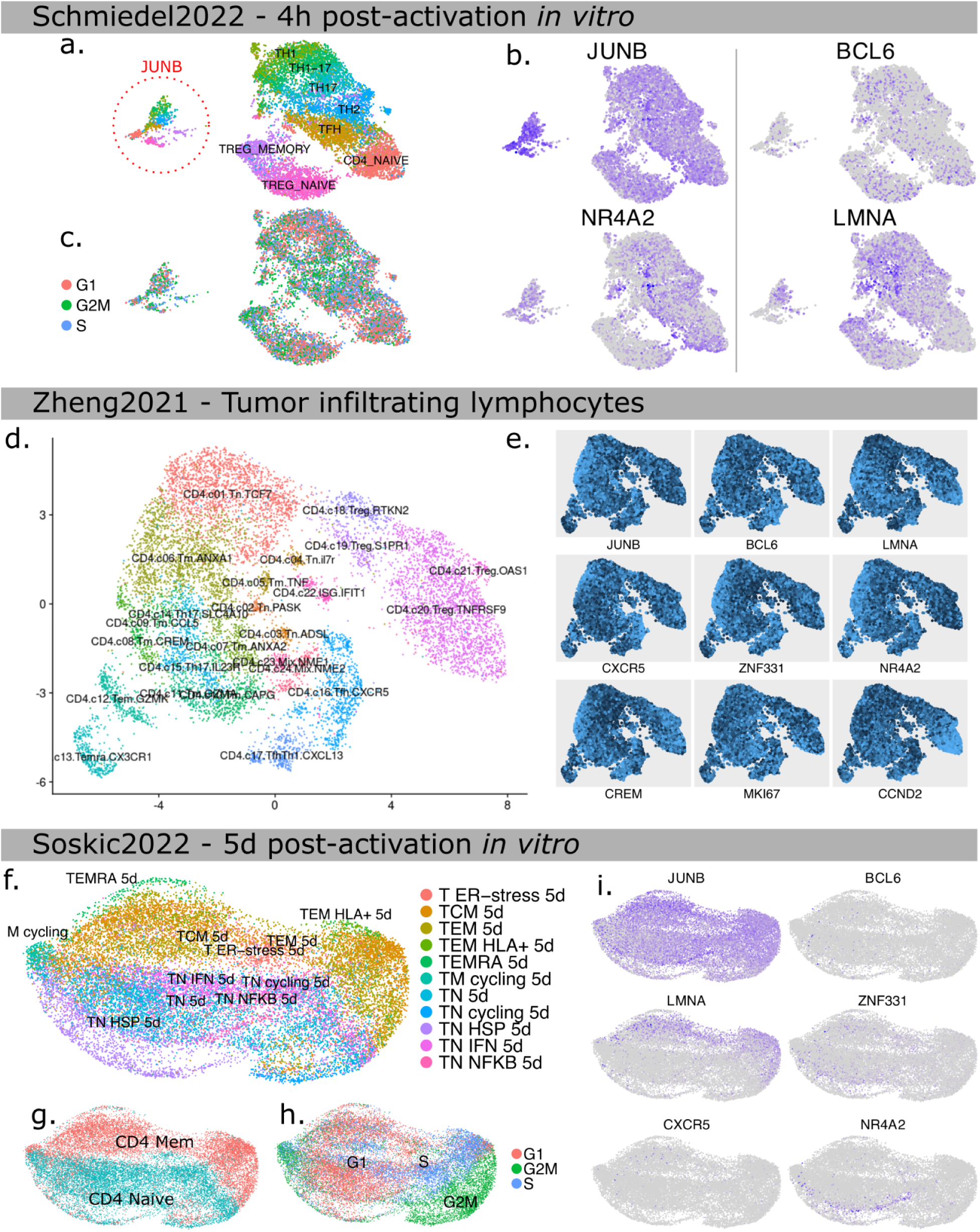
Characterization of JUNB state generality through comparison to other T cell atlases. (**a**) UMAP of in vitro activated CD4 T cell after 4h (Schmiedel2022^8^), categorized by FACS. The JUNB cluster has cells from all FACS gatings. **(b)** Marker genes discussed in text. **(c)** Cell cycle phase inferred from RNA-seq, with JUNB containing all phases. **(d)** UMAP of tumor infiltrating CD4 T lymphocytes (Zheng2021), where the distribution of JUNB is more complicated^30^. **(e)** Marker genes discussed in text, highlighting the level of co-expression to JUNB. **(f)** UMAP of Soskic2022 in vitro CD4 T cells, at 5 days after activation^31^. **(g)** UMAP colored by origin (memory or naive T cells), showing a strong separation. **(h)** Cell cycle phase inferred from RNA-seq, not correlating with JUNB expression. **(i)** Marker genes discussed in text. LMNA is in particular a good marker for memory origin CD4 T cells.

**Supplemental Figure 4:**
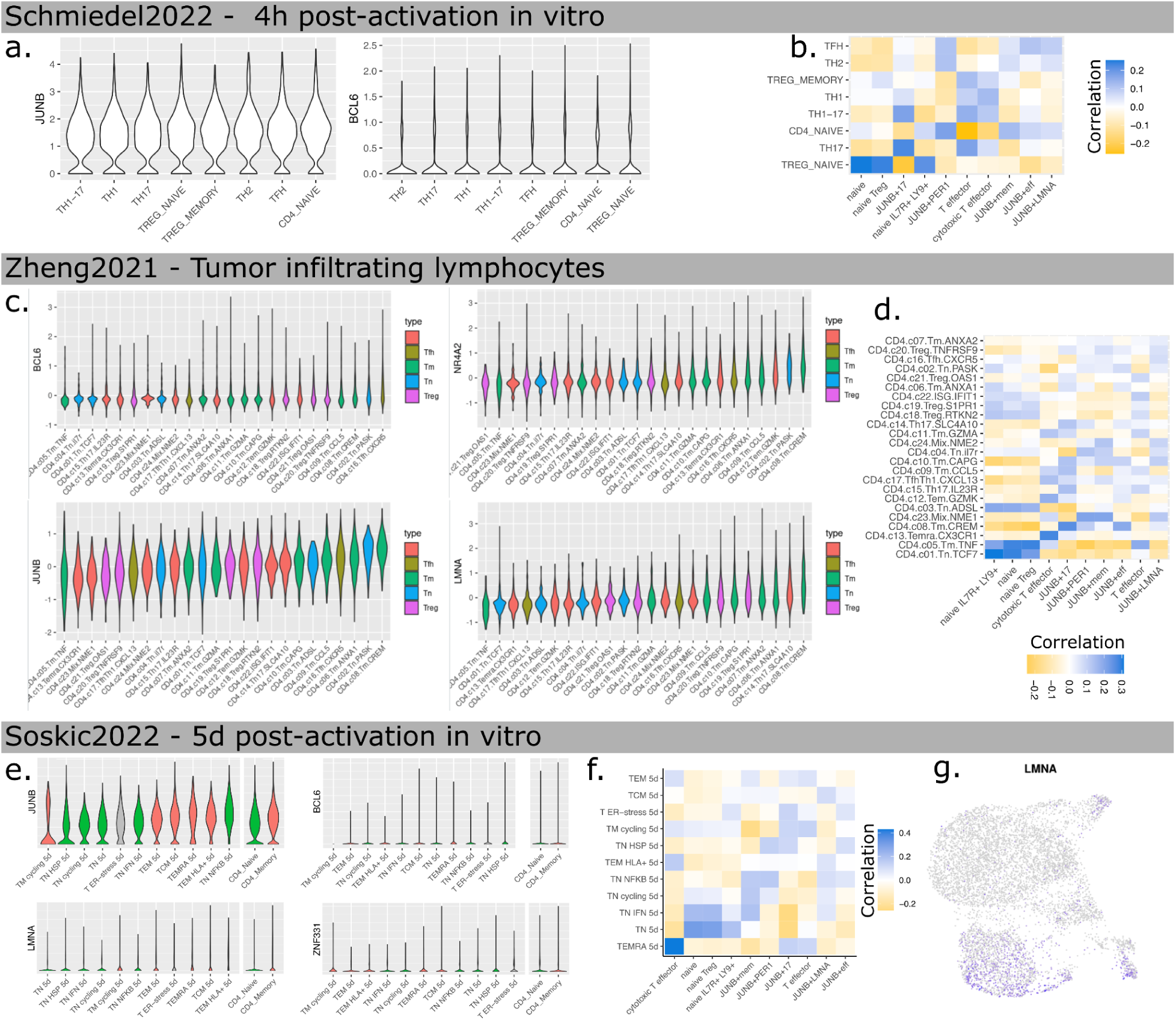
Characterization of JUNB state generality through comparison to other T cell atlases. (**a**) Expression of key genes in Schmiedel2022^8^ clusters, ordered by average level. **(b)** Genome-wide similarity of Schmiedel2022 clusters to our clusters. **(c)** Expression of key genes in Zheng2021 TIL clusters, ordered by average level. **(d)** Genome-wide similarity of Zheng2021 clusters to our clusters. **(e)** Expression of key genes in Soskic2022 clusters, ordered by average level. **(e)** Genome-wide similarity of Soskic2022 clusters to our clusters. **(f)** Genome-wide similarity of Soskic2022 clusters to our clusters. **(g)** Expression pattern of LMNA in our atlas.

**Supplemental Figure 5:**
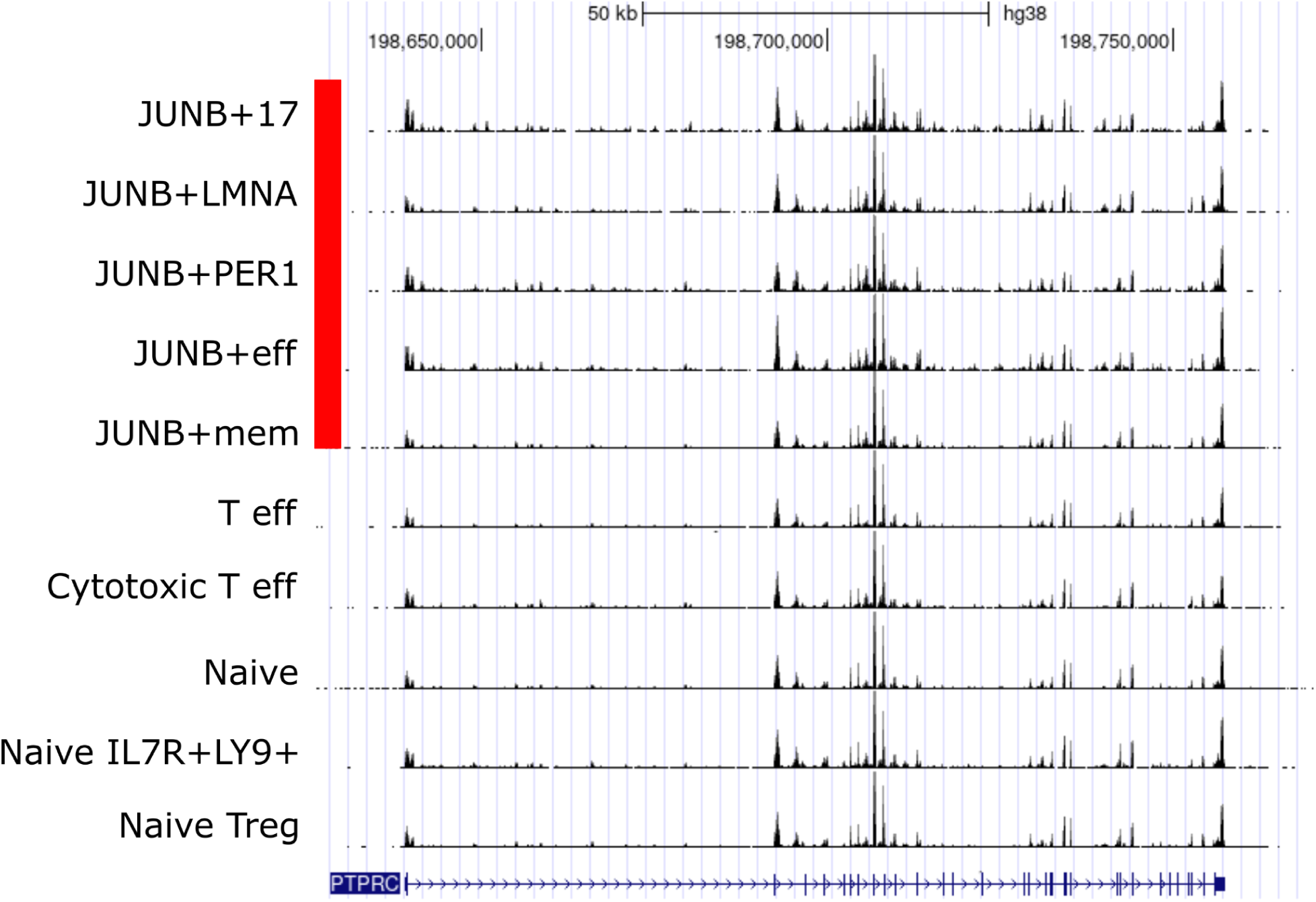
CD45 isoform analysis does not reveal particular memory T cells. Pileup of our 10x chromium RNA-seq data, over PTPRC (CD45), across all clusters in our atlas. We are unable to pinpoint any CD45RA cells using this analysis, but are also unable to conclusively rule out their presence.

We also investigated JUNB in tumor infiltrating lymphycytes^30^ (TILs, Zheng2021), reusing their cell type annotation (Figure 3d). In line with Schmiedel2022, BCL6 and JUNB expression, as well as Tfh marker CXCR5, are not isolated to the designated Tfh cluster (Figure 4e). Rather, they do not respect cluster boundaries, and are enriched in for example Tn.PASK, Tm.CREM, Tem.GZMK. This is also supported by genome-wide comparison (Figure S4d).

**Figure 4.**
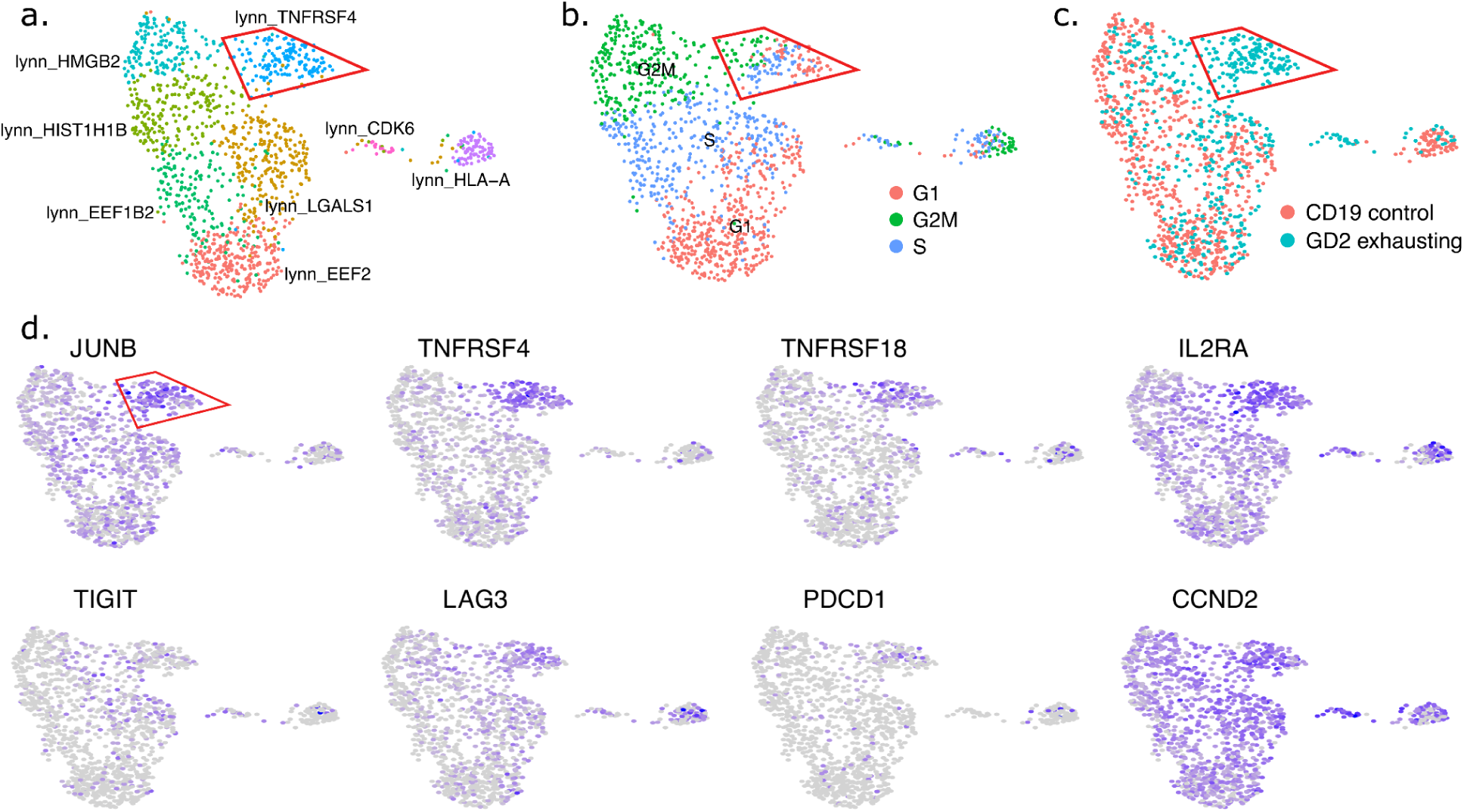
JUNB denotes a state unique to exhausting CAR T cells. (**a**) UMAP of two types of CD4 CAR T cells, one becoming exhausted due to tonic signaling. Clusters are named after their top marker gene. **(b)** Cell cycle state inferred from RNA-seq data. **(c)** Type of CAR T cell **(d)** Expression of selected genes.

**Supplemental figure 6:**
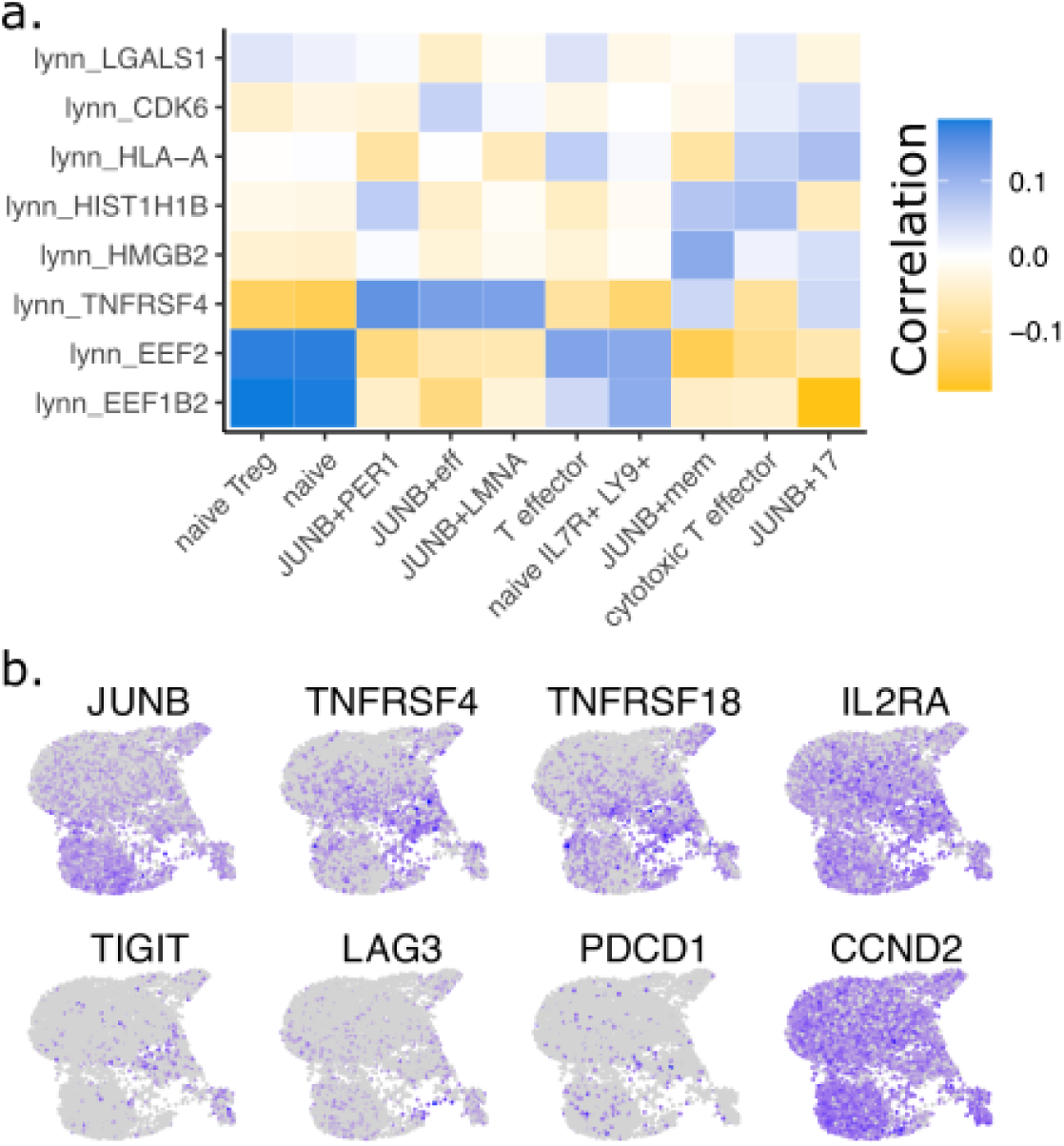
JUNB denotes a state unique to exhausting CAR T cells. (**a**) Similarity between Lynn2019 clusters and our clusters. **(b)** Genes discussed for Lynn2019, on UMAP of our atlas, for comparison.

We noted that JUNB is especially high in several TIL memory T cell clusters (Figure S4). To further investigate the potential link to memory T cells, we reanalyzed a transcriptomic *in vitro* atlas that included both naive and memory naive T cells (Soskic2022, Figure 4f). Here, the naive *vs* memory origin makes up a significant axis (Figure 3g) along with the cell cycle (Figure 3h). JUNB is elevated in memory T cells while BCL6 is not (Figure S4e). Differential expression analysis further revealed LMNA (Lamin A) to be a good marker for memory T cells (Figure 3i). Furthermore, it is up in the Soskic2022 JUNB+ cluster (and memory Treg), and in our JUNB+ cluster – especially the subset JUNB+LMNA+ (Figure 2e). However, qualitative isoform analysis of CD45 (Figure S5) does not show any clear difference between the clusters. In lack of another explanation of this cluster, we propose that this subset could be a link from naive to memory T cells, as suggested by JUNB and LMNA from the Schmiedel2022 and Soskic2022 datasets. Further experiments are however required to determine if LMNA+ is an intermediate state toward memory formation, or if it is due to an impurity from the negative selection of initial T cells.

### c-JUN and JUNB drive a proliferative and exhausting state in CAR T cells

The centrality of JUNB suggests that it might be the driving genes of the JUNB state, rather than merely being a marker for it. We thus explored datasets that would tell about causation, rather than just correlation. Overexpression of the related gene, c-JUN, in CAR T cells, has been shown to induce exhaustion resistance^32^. They also found the c-JUN-OE T cells to be more commonly in G2/M and S phase, similar to our JUNB+ cluster. But they also found other clusters affected, e.g, higher IL7R expression, and higher CD38 and IL2RA.

We reanalyzed *in vitro* single-cell data of CD19 vs GD2 CAR T cells (Lynn2019), where the latter are exhausted due to tonic signalling. We found a JUNB cluster that is present among the exhausted T cells (Figures 4a-c). It is high in effector markers IL2RA, TNFRSF4 and TNFRSF18 (Figure 4d), but also the exhaustion markers LAG3 and PDCD1. The other exhaustion marker TIGIT is elevated in a nearby cluster that (in a pseudotime sense) potentially could be a prior state. In our atlas, these markers are not expressed anywhere, plausibly because we sampled on day 5, while Lynn2019 is day 10. A genome-wide comparison shows that our JUNB+ are the most similar cluster to the Lynn2019 exhausted state (Figure S6a). TSHZ2, LMNA and NR4A2 (but not PER1) are some JUNB+ subset markers that are also enriched in the Lynn2019 JUNB cluster, suggesting these to be downstream of JUNB. Similar to the previous datasets, JUNB does not denote a certain cell cycle phase (Figure 4c). Instead, our analysis suggests it to best be thought of as an epigenetic state that favours proliferation, but which finally results in exhaustion.

### Nando, a new tool for multiome gene regulatory network analysis

The analysis so far has focused on which cell states exist, but their function and operation has only been inferred indirectly by comparison to other different environments, and different time points. However, genetic reprogramming also requires knowledge of the driver genes, and thus mechanism. The additional ATAC-seq readout enabled us to investigate the enhancer usage across cells through the Pando framework^7^. In short, by doing a motif analysis of enhancer regions, and assuming that enhancers regulate nearby genes, we can use machine learning to predict gene expression based on upstream transcription factors. The output is a function of how upstream TFs together co-regulate a gene (Figure 5a-b).

**Figure 5.**
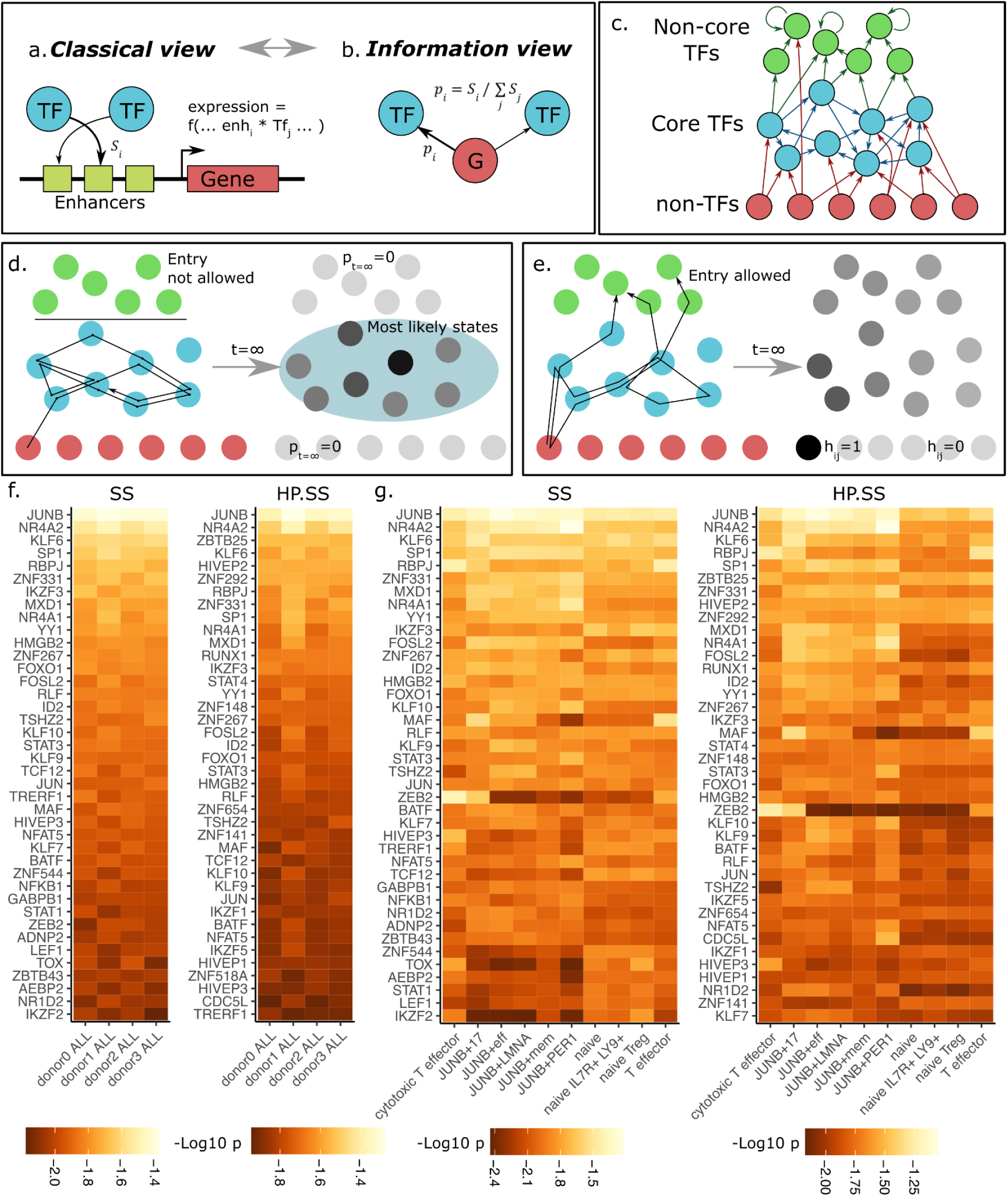
A new machine learning method is able to identify cell state driver genes. (**a**) Gene expression is predicted using machine learning over the expression of TFs and putative binding sites. **(b)** A markov chain representing the flow of information is generated based on Shapley scores. **(c)** The structure of the resulting network. **(d)** Steady state distribution showing TF centrality results from an infinite random walk. **(e)** Hitting probabilities is an alternative measure of TF importance when the random walk is allowed to get stuck among non-core TFs. **(f)** Stationary distribution, indicating the most central TFs, and Proximity (hitting probability) of upstream TFs to the irreducible network. **(g)** Centrality and hitting probability of top TFs, divided by cell type.

**Supplemental Figure 7.**
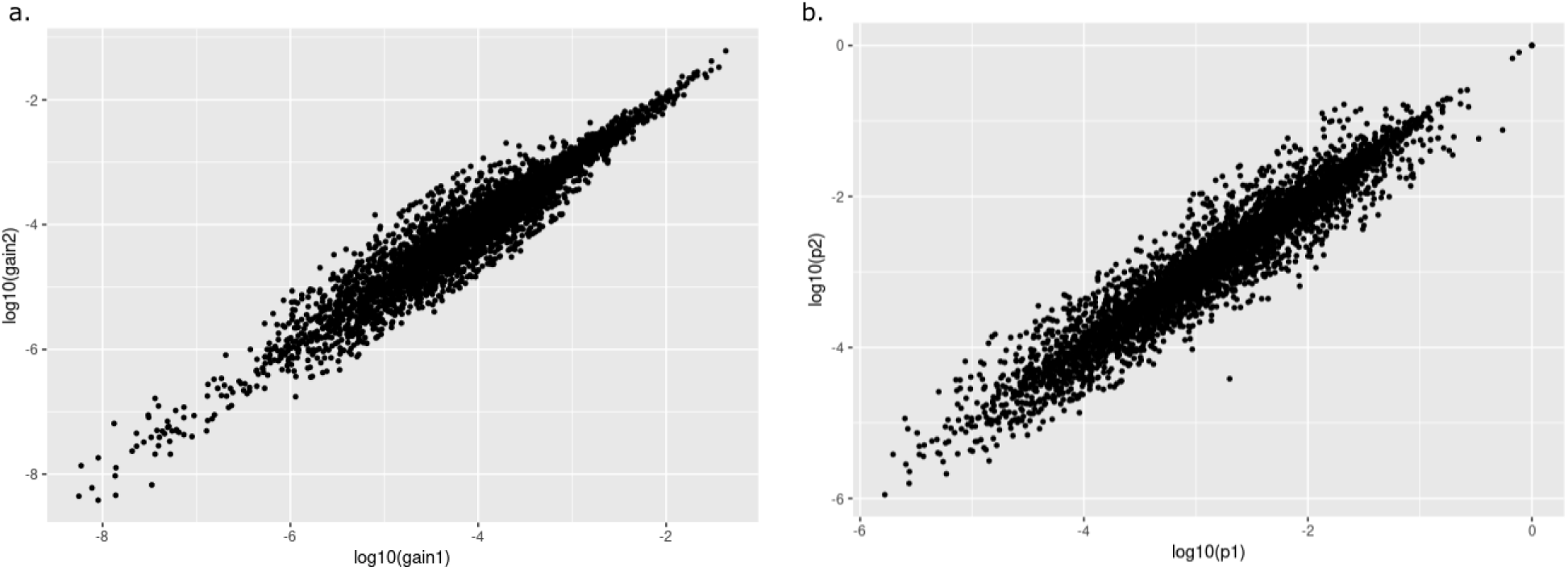
Reproducibility of network weights. A comparison of gains **(a)** and transition probabilities **(b)** over all cells, donor0 and donor1, show a high level of reproducibility even on a log scale.

An open question is how to best interpret and analyze the fitted functions. Pando currently only outputs the “gain”, i.e., how much each gene globally contributes. However, while multiple TFs binding to the enhancers of a gene, one can assume that only a subset of these TFs are actively regulating a gene in a given context (i.e. in one cell type, certain TFs are rate limiting, while other TFs mainly provide the baseline expression level). To be able to study regulation at different resolutions, we modified Pando to export the functions, and developed a new framework Nando that uses Shapley scores^33^ to interpret them. This explainable AI (XAI) approach enables efficient use of available data, as information of regulation is shared across all cells, i.e., a gene is assumed to be regulated in a roughly similar manner. This results in more stable models, and we confirm that the weights overall tend to agree across biological replicates (Figure S7).

The fitted models only predict direct interactions. To be able to study global regulation, we convert the Shapley scores into the jumping probabilities of a markov chain model (MCM)^34^ (Figure 6b). More precisely, we define these probabilities as normalized Shapley scores such that from a gene G, a jump is more likely toward a TF if it has higher regulatory potential. Thus the markov chain models where the information about the regulatory state of G most likely *came from*, and the directionality of the network (Figure 5b) is the opposite of how regulation is classically shown (Figure 5a).

**Figure 6.**
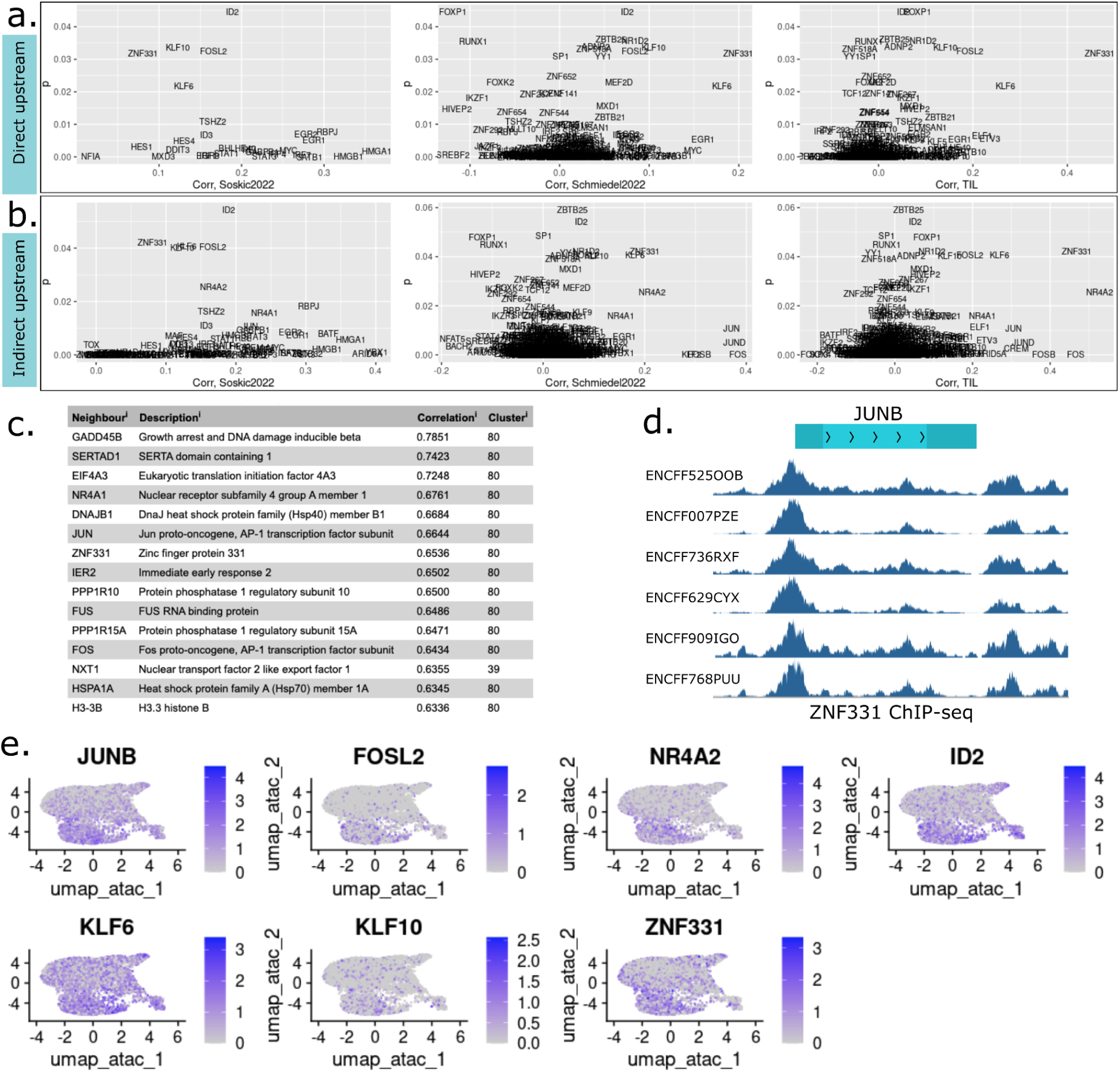
Network analysis identifies putative upstream genes of JUNB. (**a**) Candidate upstream regulators of JUNB, based on transition probability in our atlas vs correlation with genes in other atlases. Genes that are correlated also in datasets have a higher chance of being generally upstream. ZNF331 is implicated in all datasets. **(b)** Candidate upstream regulators of JUNB, based on centrality in our atlas vs correlation with genes in other atlases. **(c)** Human protein atlas closest neighbours from single cell correlation analysis show that JUNB is generally co-expressed with ZNF331 across many cell types. **(d)** HepG2 ZNF331 ChIP-seq data further shows that ZNF331 might bind near JUNB. **(e)** Some putative upstream genes, including ZNF331, plotted on UMAP of our T cell data, showing their level of co-expression with JUNB.

The resulting network consists of three types of genes based on their connectivity (Figure 4c), which affects the mathematical structure of the problem: (1) Non-TFs which have no incoming connections. (2) Core TFs connected in such a way that from any TF, a random walk is possible to any other cre TF; these make up an irreducible subgraph. (3) TFs from which it is not possible to return to the core TF subgraph. Multiple core TF subgraphs can exist but we only consider the largest such subgraph, which contains most TFs.

Markov chain theory lets us study the random walk, and thus the regulatory relationship between genes. If a walk continues forever, and starts from either core TFs or non-TFs, and never enters non-core TFs, it can most likely be found at a certain TF with probability *p*_*t*=∞_ (Figure 5d). This value is independent of the starting position and we envisage that this quantity represents how “central” or important a TF is to the overall regulation under near steady-state conditions, i.e., when a cell is not about to change its state.

The presence of non-core TFs however causes the random walk to end up stuck (“absorbed”) in some non-core TF after some time (Figure 5e). The probability of which non-core TF depends on the starting position. To mathematically be able to measure how these TFs influence regulation, we also compute the hitting probability ℎ*_ij_* which is how likely it is for a random walk from gene i to ever pass by gene j. To quantify in particular how it influences the core network, we also quantify ℎ*_j_*, the probability to pass by gene j starting from the steady state distribution.

### Explainable machine learning identifies general and cell type specific regulators

Using steady state and hitting probability analysis, we computed the globally most central TFs for each donor (Figure 5f), showing overall reproducibility. The top 10 TFs are JUNB, NR4A2, KLF6, SP1, RBPJ, ZNF331, IKZF3, MXD1, NR4A1 and YY1, with a rapid decay in centrality. TSHZ2, a member of the potential memory JUNB+LMNA, of unclear function, is among the most central TFs. It is poorly studied but has been linked to cell cycle via PRC1 in breast cancer^35^. Using hitting probabilities, it is possible to also include some additional top TFs, such as ZBTB25 (NFAT repressor^36^), HIVEP2 (expressed in Treg, and CRISPR screen shows important for Treg-mediated immunosupression^37^) and ZNF292.

As Nando makes it easy to compute cell type-specific networks, we also compared the top TFs for each cluster (Figure 5f,g, Supplemental file S10). We find that difference in centrality is often linked to the gene being a marker gene, as expected. While JUNB is central overall, it is enriched in JUNB+ as expected. So are also NR4A1/NR4A2, SP1, ZNF331, to name a few. ZEB2 is remarkably Teff-specific, while TOX and IKZF2 are relevant anywhere but Tfh. We also contrasted our results with ChromVAR (Figures 1e-g, S2f), which does not use the RNA-seq data in inferring regulators, and thus provides a complementary view. We expect the differences to be suggestive of non-transcriptional regulation, or competitive and collaborative binding to the same site. ChromVAR also only considers direct interaction unlike the Nando centrality measure. ChromVAR activity score agrees with Nando for, e.g, JUNB, but larger difference for ZNF331. In the latter case, Nando likely predicts increased centrality in Tfh due to increased ZNF331 expression, while accessibility of the motif remains the same. Nando suggests further regulators for other novel states, such as NR4A1 and CDC5L for JUNB+PER1; and TSHZ2 for JUNB+LMNA. The multiome nature of our atlas thus suggests relevant targets for modulating the CD4 T cell states, as needed to improve ACT.

### Network analysis identifies putative upstream genes of JUNB

To understand the origin of our JUNB cluster, we searched for genes upstream of JUNB. We compared the genes our network predicts with co-expression in three other CD4 T cell atlases, allowing us to filter out false positives (Figure 6a). KLF6, ZNF331, KLF10 and FOSL2 are in strong agreement as direct regulators. NR4A2 is also in agreement if considering indirect regulators (using hitting probability, Figure 6b). ZNF331 is broadly co-expressed (top 5), as is NR4A2, according to the Human protein atlas^38^ which also considers non-T cell data (Figure 6c).

Further evidence suggests that ZNF331 can act upstream, including the presence of ChIP-seq peaks at both 5’ and 3’ of JUNB (Figure 6d). ZNF331 OE also induces G2 cell cycle arrest in cancer^39^; assuming JUNB is related to cell cycle control, as suggested by our cluster, and previously shown^40^, then the phenotype of ZNF331 OE is also consistent with being upstream. However, we expect the regulation of JUNB to be more complex than being run by a single gene. PPI-analysis in Th17 shows that it is the 2^nd^ most interacting gene with FOSL2, and among top interactors for FOSL1^41^. Since FOSL2 is also being predicted as being upstream (Figure 6a), a feedback loop is plausible.

## Discussion

In this study we have elucidated the states of human primary CD4 T cells during *in vitro* culture and how knowledge of them might help improve ACT. The JUNB state turns out to have several interesting properties related to proliferation and T cell exhaustion, as inferred by comparison to most CD4 T cell atlases and T cell CRISPR screens performed to date. We thus deem it worthy of further attention for ACT purposes. They are also of interest for basic immunology, with the epigenetic topology suggesting a high degree of plasticity, which need other methods for validation, e.g. lineage tracing experiments.

The cluster JUNB+LMNA may also help us understand memory T cell formation better. We have informally come across the idea that Tfh cells may be related to memory, since they educate B cells and could benefit from being tested. Why LMNA is a marker for memory is also intriguing – we initially thought it simply a marker for dividing T cells, but the amount of protein also regulates T cell activation by modulating the TCR signal to the nucleus^42^. Because CD45 is difficult to analyze using 3’ or 5’ single-cell RNA-seq, we propose LMNA as a more viable memory marker, but needing more validation to ensure broad consistency.

A challenge in our data analysis is that BCL6 appears intertwined with JUNB, but JUNB is often expressed alone in other T cell datasets. Confounding factors is a problem both to statistics and machine learning in that models are unable to separate the source of the effects. Because we have analyzed the data using common single-cell methods, which rely on unsupervised machine learning (e.g. the UMAP and Leiden algorithms) for pattern detection and state definition, our study is no exception. Our choice of focusing on JUNB was ultimately motivated by qualitative comparison to other T cell atlases, enabling us to filter out many, possibly interesting observations, which did not seem to hold in general. Because other atlases are only transcriptomic (RNA-seq), while ours also have ATAC-seq data, we have extrapolated our results based on gene expression alone. Further multiome studies, under other conditions, are needed to see how general our results are. Overall, we are struck by the number of differences in gene expression patterns between conditions, not to say, as compared to mice. Thus we advise others optimizing T cell bioreactors for ACT to investigate the conditions anew using single-cell methods, rather than simply relying on existing atlases and a handful of marker genes.

To further inform ACT about relevant targets, we have developed a new R package and explainable ML tool, Nando, which can pinpoint cell state specific targets. By cross comparison to avoid context specific regulators, and looking for orthogonal evidence, our analysis is able to suggest ZNF331 as a general driver of the JUNB+ state. We believe other targets can be found in a similar manner. Our atlas and the gene regulatory network can be browsed online at http://data.henlab.org/.

## Methods

### T cell isolation, culture and preparation

The research was carried out according to The Code of Ethics of the World Medical Association (Declaration of Helsinki) and an ethical permit was obtained from the Swedish Ethical review authority (#2016/53-31). Blood samples were obtained from four human healthy male adults, with a mean age of 26 (ages between 20-38). Peripheral blood mononuclear cells (PBMCs) were isolated by gradient density centrifugation, using Ficoll-Paque PLUS (Cytiva, 17144002). Naïve CD4+ T cells were isolated using EasySep Human Naïve CD4+ T Cell Isolation Kit (Stemcell technologies), according to manufacturer’s indications.

Human naïve CD4+ T cells were seeded in tissue culturing 96 well plates, and stimulated with plate bound anti-human CD3 Antibody (clone OKT3, 5ng/mL, #317326, Biolegend) and aCD28 (clone CD28.2, 3 ng/mL, #302934, Biologened) at a density of 10^6^ cells/mL. The cells were activated and differentiated in ImmunoCult-XF T Cell Expansion Medium, for 5 days. The list of cytokines and concentrations used for each subtype can be found in Supplementary Table S1. Activation and differentiation stimuli were added at the same time.

On day 5, live cells were purified using flow cytometry. Cells were stained for viability discrimination using PI (P3566, Thermo Fisher Scientific), and were sorted for downstream purposes. Viability (92% on average) was then re-assessed by Trypan Blue staining (15250061, Thermo Fisher Scientific).

**Supplemental Table S1.** Reagents for the differentiation conditions.

|  | non-stimulated | Th0 | Th1 | Th2 | Th17 | Treg |
| --- | --- | --- | --- | --- | --- | --- |
| IL-2 |  | 50.00 | 50.00 | 50.00 |  | 50.00 |
| IL-6 |  |  |  |  | 5.0 |  |
| IL-4 |  |  |  | 5.0 |  |  |
| IL-12 |  |  | 5.0 |  |  |  |
| TGFb |  |  |  |  | 50.0 | 50.0 |
| IL23 |  |  |  |  | 25.0 |  |
| Atra |  |  |  |  |  | 50.0 |
| anti-IL4 |  |  | 50.0 |  |  |  |
| anti-Ifng |  |  |  | 50.0 |  |  |
| anti-cd28 |  | 15.0 | 15.0 | 15.0 | 15.0 | 15.0 |

### Validation of T cell differentiation using flow cytometry

Human naive CD4+ T cells from three different donors were seeded on a 96-well plate (1×10^6^ cells/ml, 0.2 ml/well) and activated in Th1/Th2/Th17/Treg-differentiating conditions as described above and harvested on day 5 post activation. Non-activated human naive CD4+ T cells were cultured for 5 days as well as an activation control. On day 5, live cells were purified using flow cytometry. Cells were stained for viability discrimination using PI (P3566, Thermo Fisher Scientific), and were sorted for downstream purposes. Viability (92% avg.) was then re-assessed by Trypan Blue staining (15250061, Thermo Fisher Scientific).

For the intracellular staining of TBX21 (Th1), GATA3 (Th2), FOXP3 (Treg), and RORC (Th17) the Foxp3/Transcription Factor Staining Buffer Set (00-5523, Thermo Fisher Scientific) was used according to the manufacturer’s instructions. Following antibodies were used for the labeling of aforementioned transcription factors: Human T-bet/TBX21 Alexa Fluor 488-conjugated Antibody (1:20, IC53851G, R&D Systems), Gata-3 Monoclonal Antibody (TWAJ), PE-Cyanine7 (1:20, 25-9966-42, Thermo Fisher Scientific), Human/Mouse ROR gamma t/RORC2/NR1F3 PE-conjugated Antibody (1:10, IC6006P, R&D Systems), and Human/Mouse/Rat FoxP3 PE-conjugated Antibody (1:10, IC8970P, R&D Systems). A BD FACSMelody Cell Sorter was subsequently used for the detection of the respective transcription factor.

For NKT detection, freshly isolated naive CD4+ T cells from 3 donors were analyzed for the presence of CD3, CD4, and Vα24-Jα18. Following antibodies were used for the extracellular staining according to the manufacturer’s protocol: PerCP/Cyanine5.5 anti-human CD3 Antibody (#300430, BioLegend), Brilliant Violet 421 anti-human CD4 Antibody (#357424, BioLegend), PE anti-human TCR Vα24-Jα18 (iNKT cell) Antibody (#342904, BioLegend). A BD FACSMelody Cell Sorter was subsequently used for the detection of CD3+/CD4+/Vα24-Jα18+ cells.

### Single-cell multiomics library preparation and sequencing

Dissociated cells were washed in ice-cold ATAC-seq resuspension buffer (RSB, 10 mM Tris pH 7.4, 10 mM NaCl, 3 mM MgCl2), spun down, and resuspended in 100 mL ATAC-seq lysis buffer (RSB plus 0.1% NP-40 and 0.1% Tween-20 (Thermo Fisher). Lysis was allowed to proceed on ice for 5 min, then 900 mL RSB was added before spinning down again and resuspending in 50 mL 1X Nuclei Resuspension Buffer (10x Genomics). To assess nuclei purity and integrity after lysis, nuclei were stained with Trypan Blue (15250061, Thermo Fisher Scientific), and DAPI (D1306, Thermo Fisher Scientific), according to manufacturer recommendation. If necessary, cell concentrations were adjusted to equal ratios prior to starting single-cell GEM emulsion droplet generation with the ATAC-seq NextGEM kit (10x Genomics). Briefly, nuclei were incubated in a transposition mix. Transposed nuclei were then loaded into a Chromium Next GEM Chip J. 9,000 nuclei were loaded per lane, with a target recovery of 5,500 (expected doublet rate 4% – 4.8%). After GEM emulsion generation, reverse transcription, cDNA amplification and multiome sc library generation occurred, according to manufacturer specifications (10x genomic CG000338 Rev A).

### Multiome single-cell data preprocessing

Library reads were aligned and aggregated using CellRanger ARC 2.0.0. Single-cell RNA-seq analysis was done using Seurat^43^, and single-cell ATAC-seq analysis using Signac^44^ following the standard pipeline. Briefly, initial QC filtering was performed on the scRNA-seq data (mtDNA content < 7.5%, 200 < genes/cell < 7000, 1000 < RNA-seq reads/cell < 25 000), and on the scATAC-seq data (1000 < reads/cell < 100 000, nucleosome signal < 2, TSS enrichment score >1). Peak calling was then performed anew on the filtered data using MACS2^45^.

Cell cycle scoring was performed using the list of CC-related genes^46^. Cells from respective donors were computationally separated using the available SNP information^47^. Telomemore was invoked to count telomeric reads in the ATAC-seq data, indicating primarily chromatin condensation level^26^.

### Cell type annotation

Cells from respective donors do not appear to aggregate in any particular clusters, indicating few donor differences (Figure S2a). Leiden clustering on ATAC-seq was primarily used for cell type annotation. For the LSI-based dimensional reduction, the number of PCs to include was informed by their correlation with sequence depth. Removing just the first component appeared sufficient. Several different resolutions were then tested for the Leiden clustering, with well-known marker genes qualitatively extracted at each step. Because of the high similarities between different effector subtypes (Teff), and their low percentage, they were kept as a single cluster. The manual annotation based on selected marker gene expression scores was aided by automatic annotation using DICE^48^, Monaco^49^, CellTypist^50^ (Immune_all_low.pkl as the reference), and HPA immune cell databases^49,51^. In more detail, the cell scores were based on co-expression of recently described mRNA markers for naive cells (CCR7, CD62L, TCF7, ID3, AQP3)^52^, Treg cells (FOXP3, IL2RA, TIGIT, IL2RA, CTLA4, TIGIT, TNFRSF4, TNFRSF18, PMCH, RGS1, STAT1, LGALS3)^52^, T follicular helper (Tfh) cells (BCL6, TBX21, ICOS, CXCR5, PDCD1, MAF)^53^. Subcluster of naive Tregs was assigned according to DICE annotation and reached the highest FOXP3 expression among all the clusters. According to the automatic annotation, cluster designed as “T effector” consists of Th1, Th17, and Treg cell signatures mostly (Figure S2e). Th1 signatures combined with cytotoxic markers (GZMA, GZMB, GNLY, IFNG, PRF1) were identified in the “cytotoxic T effector” cluster, which mostly comprised of CD3+ NK cells. Because of CD3+ nature the NK cells were re-annotated as NKT cells. Marker genes (FindAllMarkers) are provided as Supplemental file S8.

Cell type annotation was also attempted using Seurat batch integration and label transfer from previous *in vitro* data. The overlap was however poor. Our previous attempts at multiome B cell data^26^ label transfer has also failed, suggesting that data using the multiome chemistry is too different from plain scRNAseq for this approach to be viable.

### Cluster comparison between datasets

We generally had poor experience of several single cell batch integration methods. We have had similar issues comparing multiome B cell data^26^, and suspect it may partially be due to differences in cDNA generation between the 10x Genomics multiome RNA-seq *vs* regular single-cell RNA-seq kits. We thus instead only compared the average gene expression for each cluster across all atlases. To be precise, we computed the average gene expression per cluster. We then use Pearson correlation across the intersection of all highly variable genes, from each dataset.

### Comparison with Zheng2021 TILs

The processed TIL data^30^ was downloaded from https://doi.org/10.5281/zenodo.5461803. In particular, int.CD4.S35.sce.merged.rds was converted to Seurat. The average read count was calculated for each cluster (“meta.cluster”). Furthermore, key marker genes were qualitatively assessed based on UMAP location.

Zheng2021 dataset appears to have a large number of outliers that does not seem to stem from normal biological variation. To resolve this, we removed the top and bottom 5 cells before plotting. However, to maximize the contrast, we chose to rather plot the rank of gene expression, rather than the provided normalized values.

### Comparison with Soskic2022 *in vitro* CD4 T cells

We obtained and combined the processed files phase2_data_qced_cells_cellCycleScored.mtx.gz and phase2_data_qced_cells_cellCycleScored_cellMetadata.csv.gz. Subsetting for day 5 was done using metadata from stimulatedCells_highlyActiveCD4_5d_HVGs_processed.meta.csv. For speed, we then randomly subsampled the data to 20k cells. A routine Seurat analysis without clustering was then performed. Marker genes between memory and naive T cells were obtained by comparison over the “Cell_type” variable (CD4_Naive vs CD4_Memory).

### Comparison with Schmiedel2022 *in vitro* CD4 T cells

Raw FASTQ files were obtained from dbGaP phs001703.v4.p1 and processed using CellRanger 7.1.0. Cells were kept, having percent.mt < 10 and nCount_RNA > 4000. For speed, 1000 random cells were kept from each library. FACS gating information was obtained from the library metadata. A routine Seurat analysis without clustering was then performed.

### Comparison with Lynn2019 CAR T cells

Processed count matrices were downloaded from GEO (GSE136874) corresponding to single-cell gene expression in CD19-specific (GSM4060086) and GD2-specific (GSM4060087) CAR T cells on day 10 in culture. The matrices were concatenated and a routine Seurat workflow applied.

### Comparison with CRISPR screens

For the murine CD4 Treg differentiation (Foxp3+) screen (Cortez2020)^15^, we downloaded Table S1 and used the column “neg_lfc” as the plot y axis, and the current table row order for gene ranking x axis.

For the murine CD8 T cell exhaustion screen (Belk2022^16^), we downloaded Table S1 and used the column “z” from the sheet “genomewide_results”. We plotted against “avg_log2FC” from FindAllMarkers (logfc.threshold = 0.01) for a clustering where all a_Treg_* and a_Tfh_* were merged into two large clusters.

### Network construction and model interpretation

Pando^7^ (https://github.com/quadbio/Pando) was used to construct a GRN using all the cells, with parameters TF_corr=0, and XGBoost for model fitting. Thus the following equation is fitted for each gene.

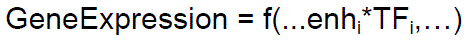

Pando default parameters were used for XGBoost^54^. We found a bug in the Pando initiate_grn. Seurat function which under certain conditions may cause gene-enhancer linkages to be scrambled. Our putative network is based on a patched version (code available at our Github).

To compute SHAP scores s_kj_, Pando was modified to (1) return the fitted models and (2) save SHAP scores for each gene and cell. Aggregated explanations S_ij_, for cell type i and gene j, were computed from all cells k as

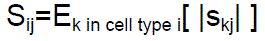

The mean and standard deviation of S_ij_ was computed across biological replicates (donors).

### Markov chain analysis

We assume the probability p_i_ of regulatory information coming from TF_i_ to be simply the normalized SHAP score 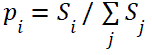. The stationary probabilities over the irreducible TFs were computed using the R package markovchain^55^, that is, it solves for probabilities ʋ in 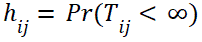. Hitting probabilities h_ij_, the probability of reaching gene j from gene i, were computed using the hittingProbabilities function. More formally, 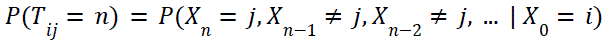 with [umath1].

### Upstream gene inference

Pearson correlation between JUNB and all other genes was computed using Rfast::correls, for all other atlases. The Lynn2019 dataset was excluded because it contains few cells, which leads to poor estimates and many false positives. We investigated top candidate upstream genes of JUNB qualitatively, expecting high correlation in all atlases.

Genes correlating with JUNB across different single cell datasets were obtained from the Humain Protein Atlas^38^ website interface (https://www.proteinatlas.org/ENSG00000171223-JUNB/single+cell+type). The pileup of ZNF331 ChIP-seq data was obtained from the online ENCODE^56^ portal (https://www.encodeproject.org/experiments/ENCSR369TCR/, corresponding to GEO entry GSE170645).

## Author contributions

I.S.M. performed the single-cell data generation, analyzed the data and wrote the first manuscript. M.S. and I.S.M. performed the validation experiments. M.S. and J.H. helped write the manuscript and analyze the data. I.M., J.T., M.F., N.B. and J.H. supervised. J.H. conceived the study. All authors have read and agreed to the published version of the manuscript.

## Supporting information

Supplemental File S8 - Markers for each cluster

Supplemental File S9 - Correlations for each TF and its motif activity

Supplemental File S10 - Gene regulatory network SS and HP.SS

## Acknowledgements

All authors have read and agreed to the published version of the manuscript.

## Supplemental Files

● Supplemental File S8: Markers for each cluster
● Supplemental File S9: Correlations for each TF and its motif activity
● Supplemental File S10: Gene regulatory network SS and HP.SS

## Data availability

The raw sequencing data has been deposited at ArrayExpress and will be made available upon acceptance. All the code is available on GitHub, including our modified version of Pando (https://github.com/henriksson-lab/Pando-fornando), Nando (https://github.com/henriksson-lab/nando) and code specific to analyzing the presented datasets (https://github.com/henriksson-lab/tcellatlas). Finally, an interactive viewer is accessible at http://data.henlab.org, along with processed count matrices, and the full Pando and Nando inputs and outputs.

